# TOR Inhibition Enhances Autophagic Flux and Immune Response in Tomato Plants Against PSTVd Infection

**DOI:** 10.1101/2024.07.11.603042

**Authors:** Samanta Silva-Valencia, Francisco Vázquez Prol, Ismael Rodrigo, Purificación Lisón, Borja Belda-Palazón

**Author notes:** Correspondence (B.B.-P.); (P.L.). These authors contributed equally to this work (Samanta Silva-Valencia, Francisco Vázquez Prol).

## Abstract

Viroids are small, non-coding RNA pathogens known for their ability to cause severe plant diseases. Despite their simple structure, viroids like Potato Spindle Tuber Viroid (PSTVd) can interfere plant cellular processes, including both transcriptional and post-transcriptional mechanisms, thereby impacting plant growth and yield. In this study, we have investigated the role of the Target Of Rapamycin (TOR) signalling pathway in modulating viroid pathogenesis in tomato plants infected with PSTVd. Our findings reveal that PSTVd infection induces the accumulation of the selective autophagy receptor NBR1, potentially inhibiting autophagic flux. Pharmacological inhibition of TOR with AZD8055 mitigated PSTVd symptomatology by reducing viroid accumulation. Furthermore, TOR inhibition promoted the recovery of autophagic flux through NBR1 and primed the plant defence response, as evidenced by enhanced expression of both the defence-related gene *PR1b* and *S5H*, a gene involved in the salicylic acid catabolism. These results suggest a novel role for TOR in regulating viroid-induced pathogenesis and highlight the potential of TOR inhibitors as tools for enhancing plant resistance against viroid infections.

## 1 Introduction

Viroids are the simplest known plant pathogens, with a genome consisting of a small (250-400 bp) circular single-stranded non-coding RNA, which is neither protected by an envelope nor encapsidated (Flores et al. 2004). Despite their simplicity, viroids exhibit great versatility. Their genome contains sufficient sequential and structural information to replicate within host cells and spread throughout the plant, producing a compatible, systemic infection that can cause severe diseases in plants, similar to those produced by viral infections (Biao 2009; Flores et al. 2014; Navarro et al. 2021). Viroids are classified into two families: *Asunviroidae* and *Pospiviroidae* (Di Serio et al. 2018, 2021). Members of the *Asunviroidae* family replicate and accumulate in chloroplasts through a rolling circle mechanism and possess hammerhead ribozyme-mediated self-cleavage, acting as catalytic RNAs. This family includes the Avocado Sunblotch Viroid (ASBVd) as its most representative member (Flores et al. 2000, 2014; Di Serio et al. 2018). *Pospiviroidae* family comprises nearly 30 known viroid species that replicate and accumulate in the nucleus. Unlike the *Asunviroidae* family, *Pospiviroidae* viroids lack self-cleaving structures (Di Serio et al. 2021). Viroid replication is considered asymmetric, following the rolling circle model similar to the *Asunviroidae* family, and beginning with a nuclear DNA-dependent RNA polymerase. The most representative member of this family is the Potato Spindle Tuber Viroid (PSTVd) (Ding et al. 2005; Flores et al. 2014; Di Serio et al. 2021).

The systemic infection produced by viroids follows a common process with several steps: entry into organelles (nucleus or chloroplasts), replication, export from the organelles, intercellular movement, entry into vascular tissue, long-distance transport through the vascular system, and subsequent entry into distal cells (Ding et al. 2005). Despite being simple organisms, viroids are infectious agents responsible for severe plant diseases, predominantly affecting higher plants and causing significant agronomic and economic impacts. This is exemplified by the diseases caused by the viroid PSTVd (Hammond and Owens 2006). PSTVd, whose genome ranges from 356 to 390 nucleotides, is a quarantine pathogen for potatoes included in the European Union’s list. Potato is the primary host and, among 156 susceptible species, 139 belong to the *Solanaceae* family, including tomato, pepper, and various ornamental species (González Arias 2014). The infection is mechanically transmitted through contact between diseased plants or via contaminated tools, among other methods. The symptoms caused by PSTVd in potato range from mild to severe, depending on the viroid strain, and can lead to a reduction in crop yield of up to 40%. Visible symptoms include clockwise twisting of the stem apex, elongated dark green leaflets, and small elongated tubers. In tomatoes, symptoms may include deformed and chlorotic leaves, shortened internodes, and stunting (Ling et al. 2012; Prol et al. 2021). PSTVd, in addition to being the reference member of the *Pospiviroidae* family, serves as an excellent model for studying viroid-host interactions due to its extensive characterization. For instance, PSTVd has been fundamental in studying viroid structure, systemic movement through the phloem, cell-to-cell movement via plasmodesmata, and developing viroid-resistant plants, among other areas (Zhu et al. 2002; Adkar-Purushothama et al. 2015, 2017). Since tomato is one of the natural hosts of PSTVd, this plant is frequently used in experimental settings due to its convenience, agronomic importance, and characteristics not present in other laboratory plants, such as its relatively short growth and maturation period of 4 to 6 weeks (Verhoeven et al. 2004). Viroids are generally considered parasites of the transcriptional machinery of organelles (nuclei or chloroplasts), unlike most plant RNA viruses, which are considered parasites of the translational machinery because they replicate in the cytoplasm and, to infect the host, must express the proteins encoded by their own genome (Tsagris et al. 2008). Viroid infection has been observed to affect the expression of genes encoding ribosomal proteins as well as proteins related to ribosome metabolism and biogenesis (Lisón et al. 2013; Góra-Sochacka et al. 2019). In addition to alterations at the transcriptional level, it has been demonstrated that viroids can affect plant cells post-transcriptionally, causing ribosomal stress in tomato plants (Cottilli et al. 2019). Specifically, citrus exocortis viroid (CEVd) was observed to cause a global defect in the translational process due, in part, to its interference with ribosome biogenesis that occurs in the nucleolus (Cottilli et al. 2019).

Stress, such as that caused by pathogen attack, affects carbon assimilation processes and ATP production, resulting in decreased energy levels (Biswal et al. 2011). Therefore, the involvement of the master energy regulator Target Of Rapamycin (TOR) in the response to such stresses is expected. Indeed, numerous studies highlight the importance of the regulation of this kinase in balancing plant growth and the defence response against pathogens (Margalha et al. 2019). During pathogen attack, TOR activity would enhance growth at the expense of the defensive response, increasing susceptibility. As a counterattack mechanism, the plant’s immune response generally involves the repression of TOR and thus diverting metabolic resources to a more efficient defensive response (Mugume et al. 2020). Consequently, plants with loss and gain-of-function of TOR tend to be more resistant and susceptible, respectively, to pathogen attack (Margalha et al. 2019).

It is important to highlight that several pathogens are capable of interfering with the action of TOR for their own benefit by activating it to promote infection. Various studies have described the interaction of viral effectors with TOR. For instance, the TAV effector protein of the Cauliflower Mosaic Virus (CAMV) can bind to and promote TOR activity, thereby aiding viral success (Schepetilnikov et al. 2011). Consequently, plants treated with specific TOR inhibitors are more resistant to CAMV infection because they lose the ability to promote polycistronic translation, thus abolishing viral replication (Schepetilnikov et al. 2011). Similarly, the inhibition of TOR by AZD8055 hinders infection caused by the Watermelon Mosaic Virus (WMV). However, infection by the Turnip Mosaic Virus (TuMV) is not affected by this inhibition, and the plants remain susceptible, suggesting that the requirement for TOR-mediated signalling may differ depending on the type of virus (Schepetilnikov et al. 2011; Ouibrahim et al. 2015).

Similar to plant-virus interactions, the activity of TOR upon bacterial infections is associated with increased susceptibility. It has been observed that TOR can partially suppress the immune response during the rice infection by *Xanthomonas oryzae* pv. *oryzae*, by neutralizing the signaling of defence hormones such as salicylic acid (SA) and jasmonic acid (JA) (De Vleesschauwer et al. 2018). Similarly, TOR inhibition in tomato plants has been shown to prime defence against several pathogens in a SA-dependent manner, including *Botrytis cinerea* (Bc), *Alternaria alternata*, and *Xanthomonas euvesicatoria* (Marash et al. 2022). In these cases, TOR inhibition induces the expression of defence-related genes, including the *PATHOGENESIS-RELATED PROTEIN 1b* (*PR1b*), a classic defence marker gene (De Vleesschauwer et al. 2018; Marash et al. 2022).

One of the key cellular processes for plant resistance and survival against pathogen attacks is autophagy (Wang et al. 2018). Autophagy is a conserved catabolic process in which cellular components such as macromolecules, damaged organelles, or toxic agents are degraded in lytic vacuoles for eventual reuse. This process is crucial during development and for maintaining cellular homeostasis under basal conditions, but it becomes particularly important under stress conditions, where it is strongly induced (Yang and Bassham 2015; Galluzzi et al. 2017).

In essence, autophagy involves the formation of a double-membrane vesicle, the autophagosome, which originates from the phagophore that surrounds and engulfs the cytoplasmic material to be degraded, transporting it to the vacuole (Marshall and Vierstra 2018). The intricate process of autophagosome initiation and maturation relies on the coordinated activity of a conserved group of AUTOPHAGY-RELATED (ATG) proteins. Within these proteins, the ATG8 family has emerged as pivotal in both the formation of autophagosomes and the recruitment of cargo. They are anchored as conjugates with the membrane lipid phosphatidylethanolamine (PE) on the expanding phagophore, enabling selectivity by interacting with a wide range of autophagic receptors and adaptors (Kushwaha et al. 2019). Autophagy can be non-selective (bulk) or selective, with the latter targeting specific components for degradation. Selective autophagy plays a crucial role in the plant immune response, relying on the lipidated ATG8 protein for specificity through interactions with various autophagic receptors known as selective autophagy receptors (Stephani and Dagdas 2020; Leong et al. 2022). One well-characterized selective autophagy receptor in plants is NEIGHBOR OF BRCA1 gene 1 (NBR1), which is essential for mediating selective autophagy-driven plant immunity by facilitating the degradation of intracellular pathogens (Svenning et al. 2011; Leong et al. 2022). In contrast to *Arabidopsis thaliana*, which harbors a single *NBR1* gene, tomato possesses two distinct *NBR1* genes: *NBR1a* (Sl03g112230) and *NBR1b* (Sl06g071770). Both genes exhibit similar intron-exon structures, resembling that of *A. thaliana NBR1*. Tomato *NBR1a* and *NBR1b* encode proteins of 864 and 738 amino acids, respectively, sharing approximately 50% sequence identity with each other and with *A. thaliana* NBR1. Like *A. thaliana* NBR1, both tomato NBR1a and NBR1b contain two highly conserved ubiquitin-associated (UBA) domains and a WxxI motif for ATG8 interaction at their respective C-termini (Zhou et al. 2014). While both NBR1a and NRB1b have been described to play important roles during heat and cold stress in tomato (Zhou et al. 2014; Chen et al. 2023), NRB1a, but not NRB1b, has been involved in root-knot nematode (RKN)-induced selective autophagy promoting tomato resistance (Yang et al. 2024).

In metazoans, it is well established that selective autophagy contributes to immune defence against viruses by actively degrading viral particles or specific proteins. This process is referred to as xenophagy (Sharma et al. 2018; Wang and Li 2020). In contrast to animals, our understanding of the roles of autophagy during plant virus infection remains still limited. Nevertheless, several studies have started to reveal the mechanisms of autophagy that are involved in plant host immunity and viral pathogenesis (Kushwaha et al. 2019; Yang and Liu 2022). Autophagy acts as a critical defence mechanism against plant DNA viruses, including geminiviruses, where it is activated during infection. For instance, the βC1 virulence factor from Cotton Leaf Curl Multan Virus (CLCuMuV) is targeted by autophagy in *Nicotiana benthamiana* through interaction with ATG8, and silencing of ATG7 and ATG5 genes enhances viral infection, highlighting autophagy’s antiviral role (Haxim et al. 2017). Additionally, βC1 functions as the first known plant viral activator of autophagy by disrupting the ATG3-glyceraldehyde-3-phosphate dehydrogenase interaction (Han et al. 2015; Ismayil et al. 2020). Similarly, other geminivirus proteins like C1 from TLCYnV are degraded via autophagy in *N. benthamiana*, mediated by interaction with ATG8h (Li et al. 2020b). Furthermore, the infection of the double-stranded DNA CaMV is inhibited by selective autophagy in *A. thaliana* (Hafrén et al. 2017). In this case, the autophagy receptor protein NBR1 targets non-assembled and virus particle-forming capsid proteins for degradation via autophagy, thereby limiting CaMV infection (Hafrén et al. 2017). All these facts highlight the role of selective autophagy in limiting viral DNA accumulation in plants. Beyond DNA viruses, autophagy also plays antiviral roles during positive-strand RNA virus infections, such as the Turnip Mosaic Virus (TuMV), where it targets viral proteins like NIb through interactions with Beclin1/ATG6 and NBR1 in *A. thaliana* (Hafrén et al. 2018). Conversely, some viruses exploit autophagy for their replication (Li et al. 2020a), while others block autophagy using viral factors to evade plant antiviral defences, highlighting the complex interplay between autophagy and viral infection in plants (Yang and Liu 2022).

Both abiotic and biotic stresses cause metabolic and energetic changes that interact with autophagy mechanisms to maintain cellular homeostasis. Adequate modulation of energy and nutrient flow is essential for plants to cope with unfavorable conditions. Therefore, the function of TOR is crucial for autophagy regulation (Wang et al. 2018; Mugume et al. 2020). Although the precise mechanisms of this regulation are mostly unknown in plants, it has been shown that TOR is a negative regulator of autophagy. Under optimal growth conditions, when the supply of nutrients and energy is normal, autophagy is maintained at low basal levels by the action of TOR, which phosphorylates ATG13, this preventing autophagy pathway initiation (Puente et al. 2016; Wallot-Hieke et al. 2018).

There is still no clear evidence directly linking the regulation of the defence response to pathogen attack through autophagy and TOR. However, an increasing number of studies correlates enhanced plant resistance with TOR inhibition (Wang et al. 2018; Margalha et al. 2019; Mugume et al. 2020). Furthermore, it has been described that the inhibition of TOR activity, which stimulates autophagy, can reduce proteotoxic stress caused by ribosomopathies or alterations in ribosome biogenesis (Recasens-Alvarez et al. 2021). Hence, studying the potential involvement of TOR signalling pathway in viroid pathogenesis, which have been described to induce alterations in ribosome biogenesis (Cottilli et al. 2019), is of significant interest.

In this study, we investigate how inhibition of TOR affects viroid disease in response to PSTVd in tomato plants. We report that PSTVd infection promotes the accumulation of the selective autophagy receptor NBR1 without increasing the autophagic flux. We demonstrate that AZD8055-mediated TOR inhibition alleviates PSTVd symptomatology by reducing viroid accumulation. Furthermore, we show that upon PSTVd infection, TOR inhibition promotes the recovery of the selective autophagic flux through NBR1, and induces the expression of the *PR1b* and *S5H*, both genes involved in plant defence. Our results demonstrate a correlation between the reduction of PSTVd accumulation and its symptomatology with the augmentation of both the autophagy flux and the defence response.

## 2 Materials and Methods

### 2.1 Plant material and growth conditions

Seeds of tomato (*Solanum lycopersicum L.*) cultivar Moneymaker were sterilized with a 1:1 mixture of commercial sodium hypochlorite and distilled H_2_O, and sequential washings of 5, 10, and 15 min were performed for the total removal of hypochlorite. Plants were cultivated in pots (12 cm deep x 13 cm inner diameter) with a 1:1 mixture of peat and vermiculite and irrigated every 2 days with Hoagland nutrient solution. The tomato plants grew in a growth chamber with a photoperiod of 16 hours of light and 8 hours of darkness. Temperature and relative humidity ranged from 28 °C and 60% during the day to 24 °C and 85% at night.

### 2.2 Viroid inoculation

In the viroid infection experiments, a strain of *Agrobacterium tumefaciens* C58 (pGV2260) expressing the dimeric cDNA sequence of PSTVd-RG1 (GenBank Acc. No. U23058.1) in the binary vector pMD201t2 was used. As a negative control for the infection, the same strain of *A. tumefaciens* containing only the plasmid was employed. The infection of tomato plants was conducted by inoculating 14-day-old plants with the appropriate *A. tumefaciens* strains, according to Prol et al. 2021. Inoculation was performed by infiltrating the abaxial side of the cotyledons using 1 mL syringes without needles. The bacterial culture expressing the viroid was grown to the exponential phase (OD_600nm_ of approximately 1.0 to 2.0). Cells were harvested by a 15 min centrifugation at 3000 *g*, and resuspended in infiltration buffer (10 mM MES, 10 mM MgCl_2_ and 200 µM acetosyringone) to achieve a final OD_600nm_ of 0.1. The agrobacteria were then incubated for 2 h at room temperature with gentle shaking before inoculation. Following infiltration, the plants were maintained under the previously described growth conditions. Leaf tissue samples were collected at specified times for further analysis.

### 2.3 TOR inhibition

TOR inhibition was performed by AZD8055 (MedChemExpress, Monmouth Junction, NJ, USA) treatment through irrigation. For this purpose, 10 mM AZD8055 stock solution prepared in DMSO was dissolved in tap water to a concentration of 1 µM (200 µL of 10 mM AZ8055 in a final volume of 2 L). Irrigation of 2 L of 1 µM AZ8055 containing solution was performed every 3 days directly into the tray (50 x 40 x 7 cm^3^) containing 6 plants per tray. As a control treatment, AZD8055 was replaced with the same volume of DMSO. The initial irrigation started one day before *A. tumefaciens* inoculation and continued for 28 days.

### 2.4 RNA extraction and RT-qPCR analyses

Total RNA was extracted from leaves using Extrazol® reagent (Blirt S.A., Gdańsk, Poland). One hundred mg of plant tissue, previously ground in liquid nitrogen, were mixed with 1 mL of the reagent in a mortar. The samples were thoroughly homogenized and then incubated at room temperature for 10 min to facilitate the dissociation of ribonucleoprotein complexes. Following this incubation, the samples were centrifuged at 13,000 *g* for 10 min at 4 °C, and the resulting supernatant was carefully transferred to a new tube. To this supernatant, 200 µL of chloroform was added, the mixture was shaken vigorously and then incubated at room temperature for 2 min. Then, the samples were centrifuged again at 13,000 *g* for 15 min at 4 °C. The resulting upper aqueous phase containing RNA was transferred to a fresh tube, and 500 µL of isopropanol was added. The mixture was gently shaken and incubated at room temperature for 10 min to precipitate the RNA. Subsequently, the RNA was pelleted by centrifugation at 13,000 *g* for 10 min at 4 °C, and the supernatant was carefully discarded. The RNA pellet was washed with 1 mL of 70% ethanol, followed by centrifugation at 7,500 *g* for 5 min at 4 °C. After discarding the supernatant, any residual ethanol was allowed to evaporate for several minutes. Finally, the RNA pellet was resuspended in 44 μL of sterile Milli-Q water treated with DEPC (diethyl pyrocarbonate) and DNA contamination was eliminated using the TURBO DNA-free™ Kit (Invitrogen, Thermo Fisher Scientific, Waltham, MA, USA), according to the manufacturer’s protocol. RNA concentration was measured using a Nanodrop™ ONE spectrophotometer (Thermo Fisher Scientific, Waltham, MA, USA) and cDNA was synthesized from 1 μg of the extracted RNA using the PrimeScript RT kit, (dT)_18_ and random primers (PerfectReal Time, Takara Bio Inc., Otsu, Shiga, Japan) following the manufacturer’s instructions.

Quantitative qRT-PCR was carried out employing PyroTaq EvaGreen qPCR Mix Plus (ROX) reagent (CMB, Madrid, Spain) with 0.4 µL of cDNA and 500 nM of each primer in a final reaction volume of 10 µL. The assays were conducted in a QuantStudio 3 Real-Time PCR System, 96-well, 0.1 cm^3^ (Applied Biosystems, Foster City, CA, USA), utilizing the *ACTIN* gene as an endogenous reference marker. Relative expression levels were calculated using the 2^-ΔCt^ method, with each biological replicate of cDNA subjected to three technical replicates. Specific sequences of the oligonucleotides used are detailed in Supplemental Table 2.

### 2.5 Protein extraction and immunologic detection

Proteins were extracted from tomato leaf tissues previously ground in liquid nitrogen. One hundred mg of ground material were mixed with 200 µL of 2× Laemmli buffer (125 mM Tris-HCl pH 6.8, 4% SDS, 20% glycerol, 4% β-mercaptoethanol, and 0.050% bromophenol blue). After homogenization, samples were kept on ice for 20 min, boiled at 95 °C for 10 min to denature the proteins and tempered on ice for 5 min. Finally, samples were centrifuged at 12,000 *g* for 15 minutes, and supernatants were preserved at -20 °C until analysis.

Samples were separated by SDS-PAGE using a Mini-Protean® Tetra handcast system (Bio-Rad, Hercules, CA, USA). The stacking gel consisted of 4% acrylamide, 0.12% SDS, 124 mM Tris-HCl pH 6.8, 0.04% APS (ammonium persulfate), 0.4% TEMED. The resolving gel consisted of 10% acrylamide, 376 mM Tris-HCL pH 8.8, 0.1% SDS, 0.1% APS, 0.2% TEMED. For the ATG8 lipidation assays, 15% acrylamide resolving gels were prepared with the addition of 6 M urea. SDS-PAGE gels were run at 90-120 V in electrophoresis buffer (25 mM Tris, 192 mM glycine, 0.1 % SDS) until the front reached the end, and used for immunologic detection.

For immunoblotting, proteins were transferred to PVDF membranes after activation with 96% ethanol for 2 minutes. Transfer occurred overnight at 12-20 V and room temperature using Bio-Rad’s (Hercules, CA, USA) wet blotting transfer system. The immunodetection process for the proteins of interest included the following steps:

- Blocking: Incubation for 1 h at room temperature with blocking buffer (5% w/v non-fat dry milk in 1× TBS containing 0.05% Tween®), with agitation.
- Primary Antibody: Overnight incubation at 4°C on a rocking platform with the primary antibody (diluted as per Table 4) in blocking buffer.
- Washing: Three washes of 10 min each with 1× TBS containing 0.05% Tween®.
- Secondary Antibody: Incubation for 1 h at room temperature on a rocking platform with the appropriate secondary antibody (as per Table 4).
- Final Washing: Three washes of 10 min each with 1× TBS containing 0.05% Tween®.
- Chemiluminescent detection was performed using suitable substrates for desired sensitivity: SuperSignal™ West Pico PLUS Chemiluminescent Substrate (Thermo Fisher Scientific, Waltham, MA, USA) and Amersham™ ECL Select™ Western Blotting Detection Reagent (Cytiva, Marlborough, MA, USA). Images were captured using a ChemiDoc system (Bio-Rad, Hercules, CA, USA) equipped with a CCD camera.

The primary antibodies used were anti-NBR1 (AS14 2805A, dilution 1:4000) and anti-ATG8 (AS14 2769, dilution 1:4000) from Agrisera, Vännäs, Sweeden. The secondary antibody was anti-rabbit IgG conjugated to horseradish peroxidase (#115035146, dilution 1:15000, Jacksons ImmunoResearch, West Grove, PA, USA). For Ponceau staining membranes were incubated with 0.1% Ponceau S (w/v) in 5% acetic acid for 10 min and washed twice with 5% acetic acid. Post-acquisition image processing and band quantification was performed using ImageJ (https://imagej.net/ij/).

### 2.6 Statistical Analysis

The statistical analyses were conducted using GraphPad Prism version 8.0.2 (https://www.graphpad.com). Comparisons between two groups were performed by either Student’s *t*-test or ratio paired *t*-test. For comparisons among more than two groups, analysis of variance (ANOVA) with Tukey’s Honestly Significant Difference (HSD) post-hoc test for multiple comparisons was performed. Unless otherwise specified, the values presented are the mean results obtained from at least 3 independent biological replicates ± the standard error of the mean (SEM). A *p*-value < 0.05 was considered statistically significant for all analyses.

## 3 Results

### 3.1 PSTVd infection promotes NBR1 accumulation due to reduced autophagy flux

Selective autophagy plays a crucial role in plant immunity by targeting and degrading intracellular pathogens, including viruses (Kushwaha et al. 2019; Leong et al. 2022). In this process, NBR1 is the only xenophagy receptor identified so far in plants (Leong et al. 2022). To determine whether PSTVd infection affects NBR1-mediated selective autophagy, tomato plants were inoculated with PSTVd, and both NBR1 protein and *NBR1a* mRNA levels were analysed 28 days after inoculation (dai), when disease symptoms such as stunting and chlorosis were clearly observable (Figure 1A). It is well stablished that the detection of NBR1 protein levels, alongside the analysis of its transcript levels, provides reliable semiquantitative data on autophagic flux in plant cells (Klionsky et al. 2016). NBR1 protein and mRNA levels were significantly up-regulated in PSTVd infected plants (Figure 1B).

**Figure 1.**
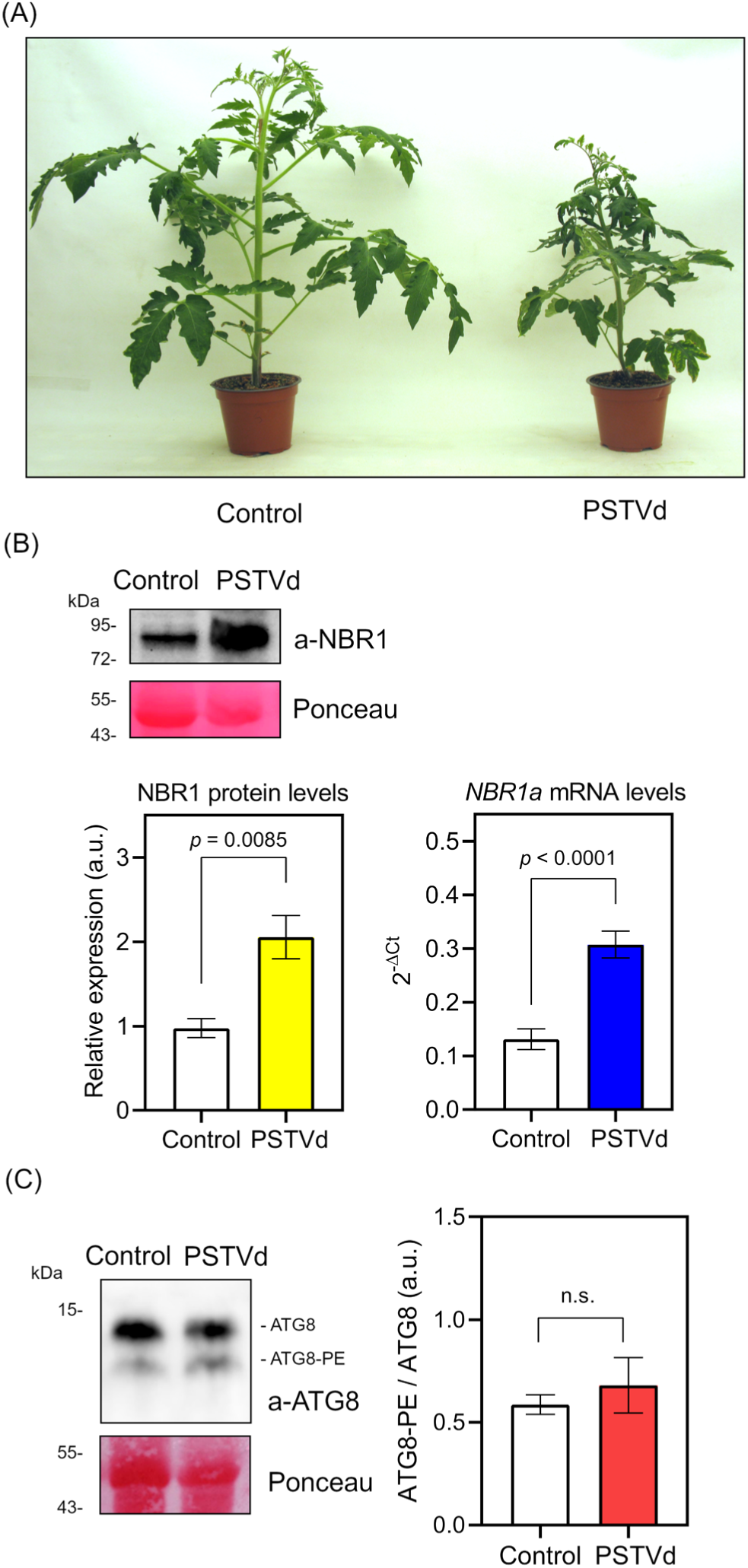
PSTVd infection promotes NBR1 accumulation due to reduced autophagy flux. (**A**) Representative pictures of non-infected (control) and PSTVd infected tomato plants at 28 dai. (**B**) Upper panels, anti-NBR1 immunoblot in control and PSTVd infected plants at 28 dai. Ponceau-S staining serves as loading control. Down left, quantification of NBR1 protein band normalized with respect to the levels of the large subunit of RuBisCO (Ponceau staining). Graph corresponds to the average of 4 biological replicates (error bars, SEM). Down right, RT-qPCR analyses of *NBR1a* gene in control and PSTVd infected plants at 28 dai. Graph correspond to the average of the relative expression levels of *NBR1a* with respect to the *ACTIN* gene (2^-ΔCt^) from 8 biological replicates (error bars, SEM). *p*-values denote statistically significant differences (two-tailed Student *t*-test). (**C**) Left, anti-ATG8 immunoblot in control and PSTVd infected plants at 28 dai, after separating protein extracts in a 6 M urea 15% SDS-PAGE. Faster-migrating bands correspond to phosphatidylethanolamine lipidated proteins (ATG8-PE). Ponceau-S staining serves as loading control. Right, quantification of the ratio between lipidated and non-lipidated ATG8 (ATG8-PE/ATG8). Graph corresponds to the average of 4 biological replicates (error bars, SEM). *p*-values denote statistically significant differences (two-tailed Student *t*-test). n.s., nonsignificant; a.u., arbitrary units.

The elevated expression of *NBR1* has been reported as a part of an induced autophagy response during plant biotic and abiotic stress (Zhou et al. 2013; Hafrén et al. 2017). Notably, the transcriptional up-regulation of *NBR1* in response to PSTVd was accompanied by a similar increase in protein accumulation (around 2-to 2.5-times in both cases). Since NBR1 itself is a substrate of the autophagy pathway (Svenning et al. 2011), our results suggest reduced NBR1 degradation and potentially impaired autophagic flux (Figure 1B). To investigate this possibility, we analysed the lipidation of ATG8 with phosphatidylethanolamine (ATG8-PE), as a hallmark of autophagy that is used as a marker to monitor autophagy flux experimentally (Reid et al. 2022). Similar levels of ATG8-PE were observed in both control and PSTVd-infected tomato plants (Figure 1C), indicating that NBR1 protein accumulation might result from reduced autophagic flux. These results underscore the role of NBR1-mediated selective autophagy in plant responses to PSTVd infection and its potential impact on disease development and plant defence mechanisms.

### 3.2 TOR inhibition reduces PSTVd accumulation and alleviates PSTVd symptomatology

The observation that PSTVd infection led to NBR1 accumulation without increasing ATG8 lipidation levels suggested that PSTVd may interfere with the autophagic process. This observation, in turn, raised the idea that TOR inhibition could enhance plant defence against PSTVd by inducing the autophagic flux (Kim et al. 2022). To verify this notion, PSTVd-infected and non-infected tomato plants were continuously treated with the TOR inhibitor AZD8055, which has been established as a *bona fide* autophagy inducer (Chresta et al. 2010; Kim et al. 2022), and analysed the PSTVd-derived symptomatology at 28 dai (Figure 2). Treatment with 1 µM AZD8055 from the moment of viroid inoculation barely affected the growth in non-infected plants, producing no severe disruption, as reported previously in *Solanaceous* species (Montané and Menand 2013; Xiong et al. 2016). Inhibition of TOR by AZD8055 caused symptomatic relief in PSTVd-infected tomato plants compared to untreated plants. While PSTVd infection led to a progressive decrease in internodal length and total plant height, TOR inhibition in infected plants partially restored plant growth (Figure 2A and B, and Supplemental Table 1). To test whether the symptomatic relief caused by TOR activity inhibition was due to a decrease in viroid accumulation, the viroid levels were analysed in these plants (Figure 2C). Inhibition of TOR by AZD8055 caused a significant reduction in the accumulation of PSTVd of tomato-infected plants. These findings collectively suggest that TOR inhibition with AZ8055 alleviates PSTVd-induced symptoms in tomato plants by reducing viroid accumulation, thus highlighting TOR’s potential role in modulating plant defence mechanisms against viroid infections.

**Figure 2.**
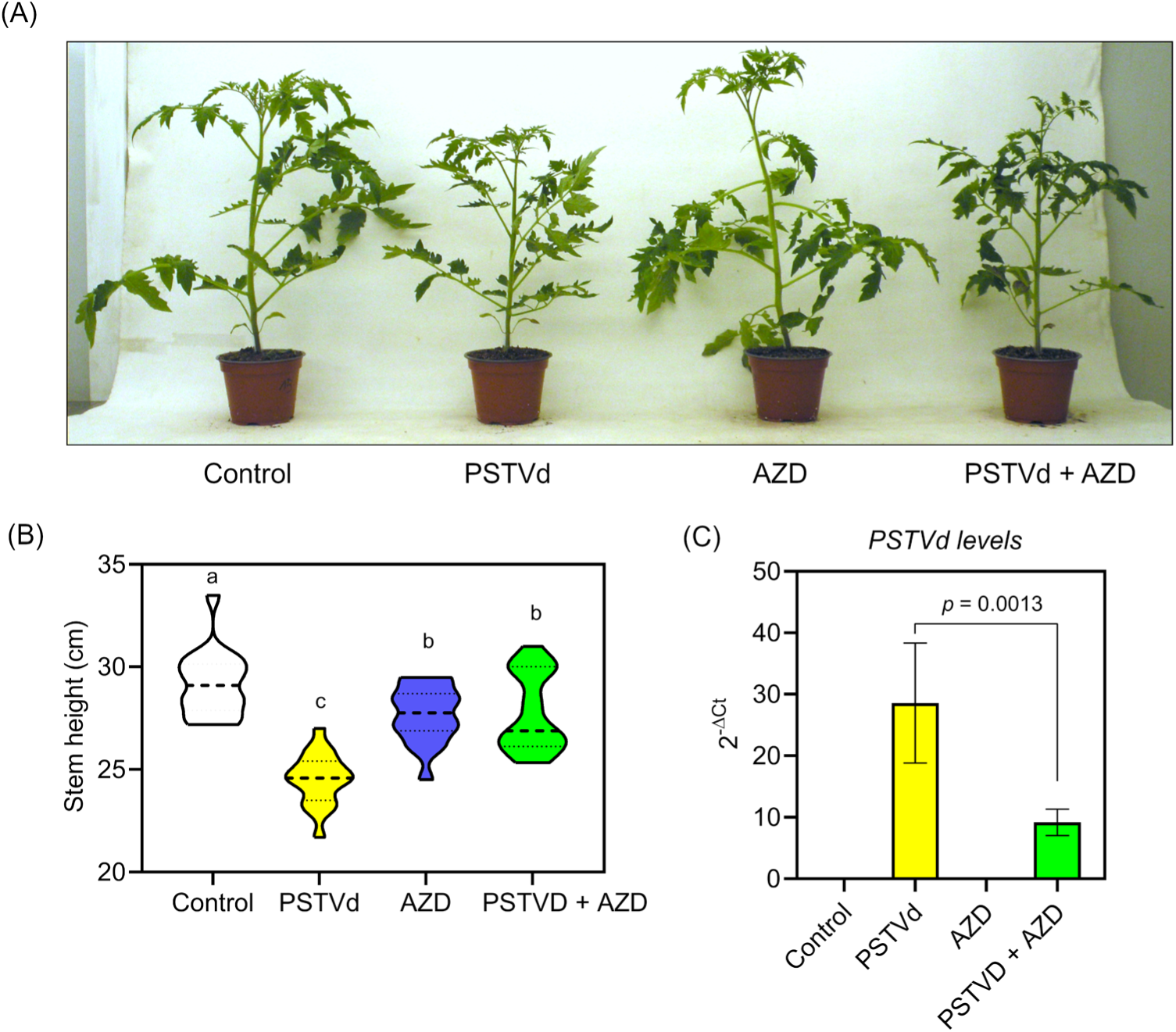
TOR inhibition reduces PSTVd accumulation, alleviating PSTVd symptomatology. (**A**) Representative pictures of non-infected (control) and PSTVd infected tomato plants at 28 dai, with or without continuous treatment with AZD8055. (**B**) Violin box plots representing height measurements of 18 plants, arranged with 6 plants per tray, across 3 independent experiments. Different letters denote statistically significant differences (*p* < 0.05, one-way ANOVA with Tukey HSD test). (**C**) RT-qPCR analyses of PSTVd levels in the specified treatments at 28 dai. Graph corresponds to the average of the relative levels of PSTVd with respect to the *ACTIN* gene (2^-ΔCt^) from 10 plants (error bars, SEM). *p*-values denote statistically significant differences (ratio paired t-test).

### 3.3 TOR inhibition in PSTVd-infected plants restores the autophagic flux

The alleviation of viroidal symptoms and reduction in PSTVd levels in infected plants by the inhibition of TOR activity raised the hypothesis of an activated autophagy. To explore this correlation, we analysed autophagic flux by assessing the accumulation of *NBR1* transcript and protein levels in both PSTVd-infected and non-infected tomato plants treated with the TOR inhibitor AZ8055 (Figure 3A and B). As expected, treatment with AZ8055 of non-infected plants induced an increase in *NBR1a* transcript levels with no significant increase in NBR1 protein levels (Figure 3A and B), as established evidence of induced autophagy (Zhou et al. 2013; Hafrén et al. 2017). Interestingly, in PSTVd-infected plants, inhibition of TOR resulted in a significant reduction in NBR1 protein accumulation, which was not accompanied by a significant decrease in transcript levels, unlike what was observed in non-treated PSTVd-infected plants where NBR1 protein levels were strongly induced to a similar extent as *NBR1a* transcript levels. These results indicate that AZD8055-mediated TOR inhibition restored the autophagic flux in PSTVd-infected tomato plants.

**Figure 3.**
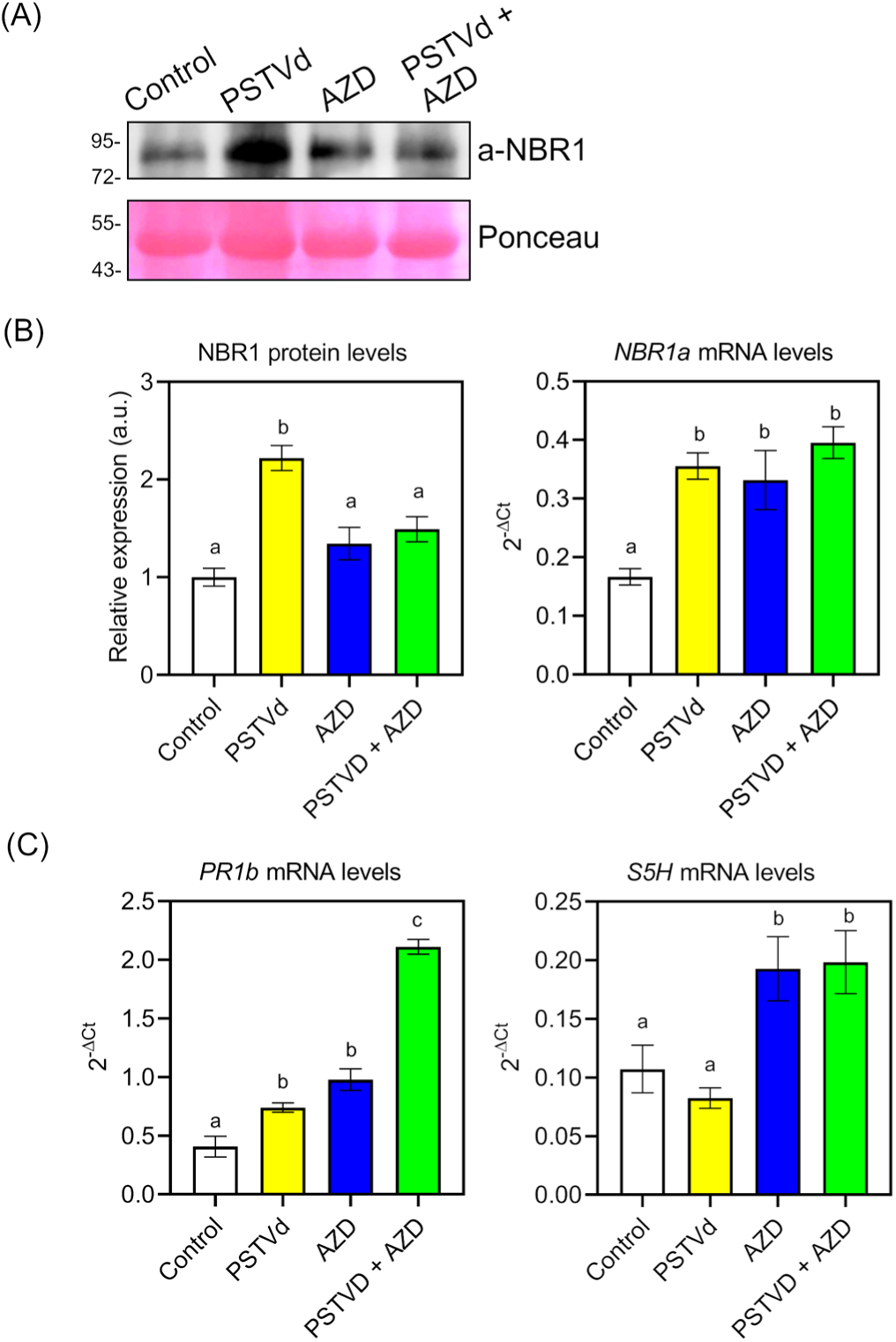
TOR inhibition restores the autophagic flux and primes defense mechanisms in PSTVd-infected plants. (**A**) Anti-NBR1 immunoblot in non-infected (control) and PSTVd infected tomato plants at 28 dai, with or without continuous treatment with AZD8055. Ponceau-S staining serves as loading control. (**B**) Left, quantification of NBR1 protein normalized with respect to the levels of the large subunit of RuBisCO (Ponceau staining). Graph corresponds to the average of 6 biological replicates (error bars, SEM). Right, RT-qPCR analyses of *NBR1a* gene for each treatment. Graph corresponds to the average of the relative expression levels of *NBR1a* with respect to the *ACTIN* gene (2^-ΔCt^) from 4 biological replicates (error bars, SEM). (**C**) RT-qPCR analyses of *PR1b* and *S5H* genes for each treatment. Graphs correspond to the average of the relative expression levels of *PR1b* and *S5H* with respect to the *ACTIN* gene (2^-ΔCt^) from 4-5 biological replicates (error bars, SEM). (**B** and **C**) Different letters denote statistically significant differences (*p* < 0.05, one-way ANOVA with Tukey HSD test).

### 3.4 TOR inhibition primes the defence response in PSTVd-infected plants

TOR inhibition enhances immunity against diverse pathogens including virus and bacteria (De Vleesschauwer et al. 2018; Mugume et al. 2020; Marash et al. 2022). To better understand the impact of TOR down-regulation on defence mechanisms triggered by PSTVd infection, we investigated the transcriptional response of the *PATHOGENESIS-RELATED PROTEIN 1b* (*PR1b*), a well-known defence marker gene induced by both systemic acquired resistance (SAR) and induced systemic resistance (ISR) (Meller Harel et al. 2014; Li et al. 2017), to AZD8055 treatment alone and in conjunction with PSTVd infection (Figure 3C). Consistent with previous reports (De Vleesschauwer et al. 2018; Prol et al. 2021; Marash et al. 2022), the expression levels of *PR1b* showed a significant increase (approximately 2-fold) in either PSTVd-infected or TOR-inhibited plants. Interestingly, continuous AZD8055 treatment of PSTVd-infected plants further enhanced the expression of *PR1b* (approximately 4-fold higher than control plants, and 2-fold higher than TOR-inhibited or PSTVd-infected plants), indicating an additive effect between PSTVd infection and TOR inhibition in priming immunity and enhancing disease resistance (Figure 3C). Moreover, the induction of *PR1b* in AZD8055 treated plants was accompanied by an induction of *S5H*, a gene involved in the SA catabolism which is induced by SA (Payá et al. 2022). These results suggest that AZD8055-mediated TOR inhibition activates the SA-mediated tomato immune response.

## 4 Discussion

Selective autophagy is a crucial cellular process that enables plants to survive and resist pathogen attacks, for which selective autophagy receptors play a vital role by recognizing intracellular pathogenic components such as viral proteins and bacterial effectors and facilitating autophagosome formation (Sharma et al. 2018; Wang et al. 2018; Leong et al. 2022). NBR1 is the only known xenophagy cargo receptor in plants (Leong et al., 2022),and while its role in virus infections has only recently started to emerge, its role in infections caused by non-coding single-stranded RNA viroids remains completely unknown. In this work, it was demonstrated that the infection of tomato plants by PSTVd induced the accumulation of NBR1 at both the transcript and protein levels (Figure 1B), suggesting that PSTVd may hinder defensive autophagy, as confirmed by the unchanged lipidation of ATG8 (Figure 1C). Similarly, the counteraction of NBR1-mediated selective autophagy has been demonstrated for positive-stranded RNA viruses. For instance, TuMV disrupts autophagic flux through the action of distinct viral proteins, leading to NBR1 accumulation (Hafrén et al. 2018). This contrasts with other DNA viruses that promote selective autophagy, such as CaMV, which induces the transcriptional up-regulation of NBR1 without simultaneous protein accumulation, indicating enhanced NBR1 turnover and increased autophagic flux (Hafrén et al. 2017). The molecular mechanisms by which PSTVd, lacking coding capacity, might interfere with the autophagy process remain to be fully elucidated.

TOR has been identified as a key regulator of autophagy (Mugume et al. 2020). Its role in plant defence against various pathogens, including viruses, bacteria and fungi, has been extensively studied, demonstrating that in most cases, inhibition of TOR can enhance plant resistance to these pathogens (Margalha et al. 2019). However, despite the acknowledged importance of TOR in regulating autophagy and its implications for plant defence responses, the evidence demonstrating a direct relationship between TOR-regulated autophagy and the defence response to pathogen attack in plants remains limited (Zvereva et al. 2016). In this work, it was hypothesized that inhibition of TOR could prime defence against PSTVd in tomato plants. The results demonstrated that blocking TOR significantly reduced PSTVd levels, leading to symptomatic relief of viroid infection in tomato plants (Figure 2A-C). This alleviation could be attributed to the restoration of autophagic activity, as indicated by the strong correlation observed between symptomatic relief, reduction in PSTVd levels, and the upregulation of transcript levels coupled with the downregulation of protein levels of NBR1 (Figure 3A and B).

TOR inhibition has been shown to enhance immunity against diverse pathogens (De Vleesschauwer et al. 2018; Marash et al. 2022). Therefore, we hypothesized that pharmacological inhibition of TOR could similarly facilitate an immune response against PSTVd. This study has demonstrated that TOR inhibition in PSTVd-infected plants induce the immune response, as evidenced by the induction of *PR1b* (Figure 3C). Previous reports have shown that both PSTVd infection and TOR inhibition individually activate *PR1b* (Prol et al. 2021; Marash et al. 2022). In contrast, TOR inhibition in PSTVd-infected plants resulted in an additive effect in *PR1b* induction, which could be attributed to either synergistic or separate individual effects. This differs from the non-additive effect observed in the immune response against Bc infection, where TOR inhibition did not further increase the expression of defence genes such as *PR1b* (Marash et al. 2022). Interestingly, SA, one of the classic defence hormones, has been shown to be antagonized by TOR (De Vleesschauwer et al. 2018). In this connection, PSTVd-infection elevated the levels of SA (Prol et al. 2021), its exogenous application has been shown to improve the resistance to PSTVd (Li et al. 2021), and plants with impaired accumulation of SA are more susceptible to CEVd (López-Gresa et al. 2016). Furthermore, SA has the ability to modulate autophagy (Yoshimoto et al. 2009; Shukla et al. 2022). Consistent with this, we have observed that AZD8055-mediated TOR inhibition also induced *S5H* (Figure 3C), a salicylic acid 5-hydroxylase that has been described to be induced by SA (Payá et al. 2022). Therefore, the activation of both *PR1b* and *S5H* by TOR inhibition appears to indicate that the observed reduction in PSTVd levels and symptoms could also be due to the activation of the tomato SA-mediated defence response. Alternatively, it has been recently shown that inhibition of TOR activity reduces proteotoxic stress caused by ribosomopathies (Recasens-Alvarez et al. 2021). We could therefore hypothesize that the symptomatic relief caused by TOR inhibition could be due not only to the activation of autophagic flux and plant defence, but also to the alleviation of ribosomopathies caused by viroids, as they have been observed to associate with ribosomes, producing alterations in their biogenesis (Cottilli et al. 2019).

The results presented in this work indicate that manipulating the growth-defence switch via TOR inhibition can prime tomato immunity and disease resistance against PSTVd, opening new perspectives on cellular mechanisms previously not considered in viroid pathogenesis. These mechanisms include the role of NBR1-mediated selective autophagy and its interaction with TOR.

## 5 Conclusions

This study explores TOR inhibitory effects on viroid disease in tomato plants infected with PSTVd. We found that PSTVd infection promotes NBR1 accumulation without increasing autophagic flux. TOR inhibition with AZD8055 alleviates PSTVd symptoms by reducing viroid levels, restoring selective autophagic flux through NBR1, and inducing defence-related protein *PR1b* expression, demonstrating the link between enhanced autophagy and reduced viroid pathogenicity.

## Author contributions

S.S.-V, F.V.P. and B.B.-P. conducted experiments. I.R. provided expertise on PSTVd signalling, actively supporting the conceptual work. B.B.-P. and P.L. conceived and designed the project. B.B.-P. directed and supervised all research activities and prepared figures. B-B.-P. wrote the manuscript. P.L. and I.R. reviewed the manuscript. All authors read and approved the final manuscript.

## Acknowledgments

We acknowledge the financial support received for the development of this work. We would also like to thank the support offered by the services of the IBMCP (Valencia, Spain).

## Data availability statement

All data supporting the findings of this study are available in both the main text and supplemental information. Additional data related to this study are available from the corresponding authors upon request.

## Reference

**Supplemental Table 1.**
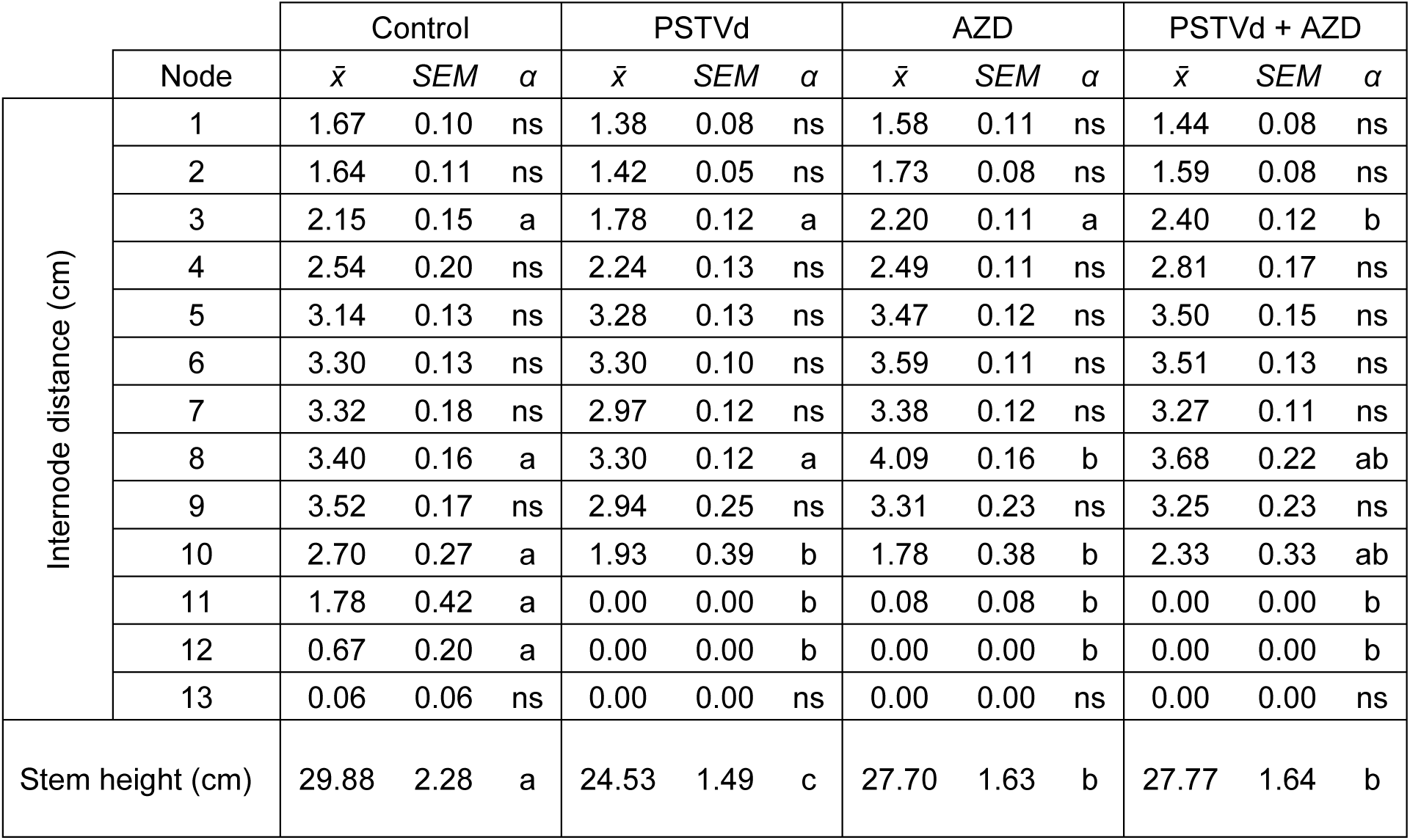
Internode distance quantification of non-infected (control) and PSTVd infected tomato plants at 28 dai, with or without continuous treatment with AZD8055. *x̂*, mean of 18 plants, arranged with 6 plants per tray, across 3 independent experiments. SEM, standard error of the mean. *α*, one-way ANOVA with Tukey HSD test. Different letters denote statistically significant differences (*p* < 0.05).

**Supplemental Table 2.**
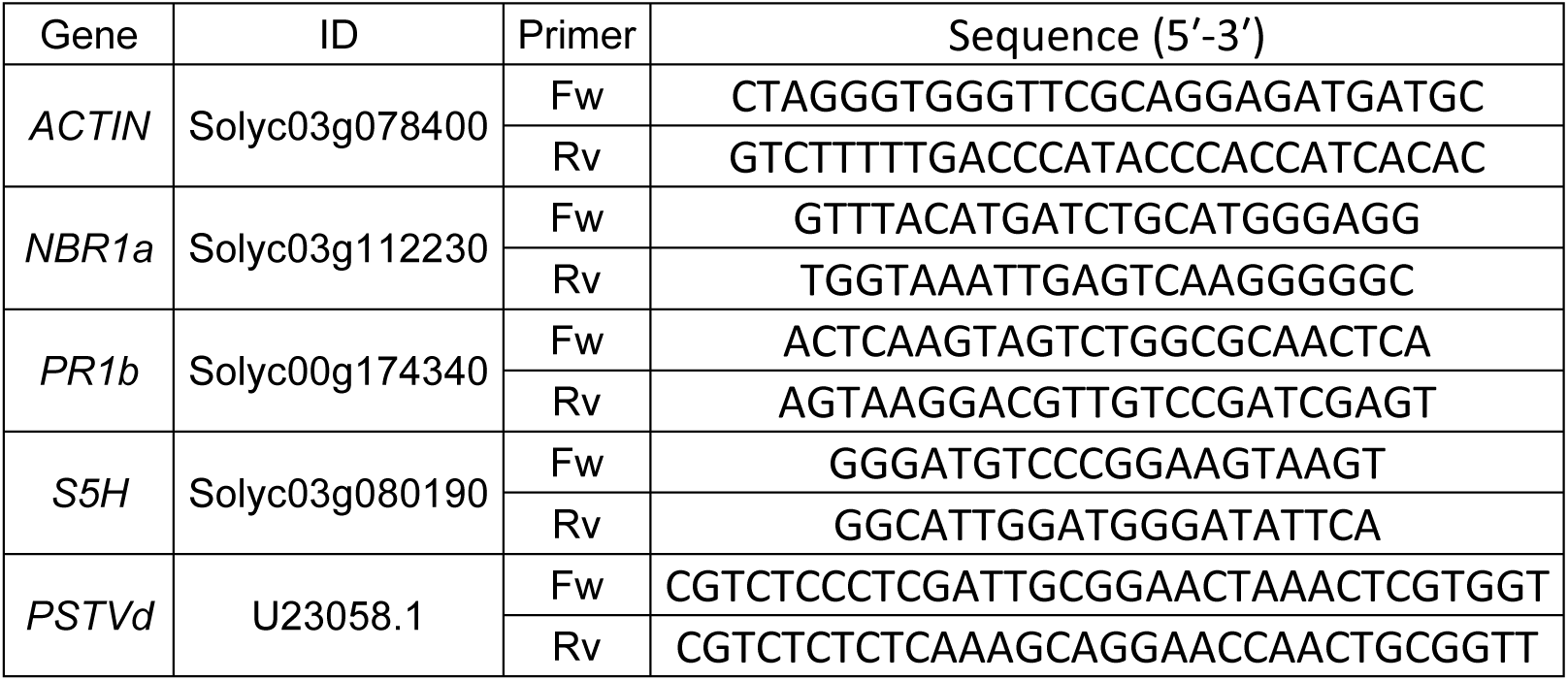
List of primers used in this work.

