## Supplemental Table 1 and 2 for "TOR Inhibition Enhances Autophagic Flux and Immune Response in Tomato Plants Against PSTVd Infection"

**Supplemental Table 1**. Internode distance quantification of non-infected (control) and PSTVd infected tomato plants at 28 dai, with or without continuous treatment with AZD8055. *x̄*, mean of 18 plants, arranged with 6 plants per tray, across 3 independent experiments. SEM, standard error of the mean. *α*, one-way ANOVA with Tukey HSD test. Different letters denote statistically significant differences (*p* < 0.05).

|  |  | Control | | | PSTVd | | | AZD | | | PSTVd + AZD | | |
| --- | --- | --- | --- | --- | --- | --- | --- | --- | --- | --- | --- | --- | --- |
|  | Node | *x̄* | *SEM* | *α* | *x̄* | *SEM* | *α* | *x̄* | *SEM* | *α* | *x̄* | *SEM* | *α* |
| Internode distance (cm) | 1 | 1.67 | 0.10 | ns | 1.38 | 0.08 | ns | 1.58 | 0.11 | ns | 1.44 | 0.08 | ns |
|  | 2 | 1.64 | 0.11 | ns | 1.42 | 0.05 | ns | 1.73 | 0.08 | ns | 1.59 | 0.08 | ns |
|  | 3 | 2.15 | 0.15 | a | 1.78 | 0.12 | a | 2.20 | 0.11 | a | 2.40 | 0.12 | b |
|  | 4 | 2.54 | 0.20 | ns | 2.24 | 0.13 | ns | 2.49 | 0.11 | ns | 2.81 | 0.17 | ns |
|  | 5 | 3.14 | 0.13 | ns | 3.28 | 0.13 | ns | 3.47 | 0.12 | ns | 3.50 | 0.15 | ns |
|  | 6 | 3.30 | 0.13 | ns | 3.30 | 0.10 | ns | 3.59 | 0.11 | ns | 3.51 | 0.13 | ns |
|  | 7 | 3.32 | 0.18 | ns | 2.97 | 0.12 | ns | 3.38 | 0.12 | ns | 3.27 | 0.11 | ns |
|  | 8 | 3.40 | 0.16 | a | 3.30 | 0.12 | a | 4.09 | 0.16 | b | 3.68 | 0.22 | ab |
|  | 9 | 3.52 | 0.17 | ns | 2.94 | 0.25 | ns | 3.31 | 0.23 | ns | 3.25 | 0.23 | ns |
|  | 10 | 2.70 | 0.27 | a | 1.93 | 0.39 | b | 1.78 | 0.38 | b | 2.33 | 0.33 | ab |
|  | 11 | 1.78 | 0.42 | a | 0.00 | 0.00 | b | 0.08 | 0.08 | b | 0.00 | 0.00 | b |
|  | 12 | 0.67 | 0.20 | a | 0.00 | 0.00 | b | 0.00 | 0.00 | b | 0.00 | 0.00 | b |
|  | 13 | 0.06 | 0.06 | ns | 0.00 | 0.00 | ns | 0.00 | 0.00 | ns | 0.00 | 0.00 | ns |
| Stem height (cm) | | 29.88 | 2.28 | a | 24.53 | 1.49 | c | 27.70 | 1.63 | b | 27.77 | 1.64 | b |

**Supplemental Table 2**. List of primers used in this work.

| Gene | ID | Primer | Sequence (5′-3′) |
| --- | --- | --- | --- |
| *ACTIN* | Solyc03g078400 | Fw | CTAGGGTGGGTTCGCAGGAGATGATGC |
|  |  | Rv | GTCTTTTTGACCCATACCCACCATCACAC |
| *NBR1a* | Solyc03g112230 | Fw | GTTTACATGATCTGCATGGGAGG |
|  |  | Rv | TGGTAAATTGAGTCAAGGGGGC |
| *PR1b* | Solyc00g174340 | Fw | ACTCAAGTAGTCTGGCGCAACTCA |
|  |  | Rv | AGTAAGGACGTTGTCCGATCGAGT |
| *S5H* | Solyc03g080190 | Fw | GGGATGTCCCGGAAGTAAGT |
|  |  | Rv | GGCATTGGATGGGATATTCA |
| *PSTVd* | U23058.1 | Fw | CGTCTCCCTCGATTGCGGAACTAAACTCGTGGT |
|  |  | Rv | CGTCTCTCTCAAAGCAGGAACCAACTGCGGTT |
